# TMX1, a disulfide oxidoreductase, is necessary for T cell function through regulation of CD3ζ

**DOI:** 10.1101/2024.09.22.614388

**Authors:** Timothy Chai, Kyle M. Loh, Irving L. Weissman

## Abstract

T cell-targeted therapies are commonly used to manage T cell hyperactivity in autoimmune disorders, graft-versus-host diseases (GVHD), and transplant rejections. However, many patients experience significant side effects or inadequate responses to current treatments, highlighting the urgent need for alternative strategies. In this study, we searched for regulators of T cells through proximity labeling with APEX2 to detect proteins interacting with CD8α, a coreceptor of the T-cell receptor (TCR). This screen revealed TMX1, an ER resident transmembrane disulfide oxidoreductase, is essential for T cell cytotoxicity and NFAT, NFκB, and AP1 signaling but not cell proliferation. TMX1 deletion decreases surface TCR expression and destabilizes CD3ζ, a subunit of TCR complex; however, overexpression of CD3ζ rescues the phenotype, suggesting that TMX1 is not required for CD3ζ function. Mechanistically, TMX1 was found to directly engage the CxxC motif of CD3δ, which has been reported to be essential for proper TCR assembly and function. We hypothesize that the loss of TMX1 interaction with CD3δ leads to impaired TCR assembly and subsequent CD3ζ destabilization. These findings identify TMX1 as a novel regulator of T-cell receptor assembly and a potential target for immunosuppressive therapy.

## Introduction

T cell hyperactivity affects millions of people worldwide^1,2^, including those with asthma^3,4^, rheumatoid arthritis^5^, type 1 diabetes ^6^, inflammatory bowel disease ^7^, multiple sclerosis^8^, among others. These diseases significantly reduce the quality of life, often exacerbated by side effects of immunosuppression or steroids ^9–17^. T cell hyperactivity poses significant risk of mortality in more severe conditions, such as graft-versus-host disease (GVHD) ^18–20^ and organ transplant rejection^21–23^. The need for effective strategies to control T cell hyperactivity is therefore vital in improving outcomes for affected individuals.

As transplants has become the mainstay treatment for end stage organ diseases and subsets of cancers, the number of organ and bone marrow transplants has increased ^24–26^. Despite advances in prophylactic treatments, GVHD develops in approximately 50% of patients undergoing bone marrow transplants ^27–30^. For those with grade II–IV acute GVHD, 30-60 % are steroid refractory ^31,32^ and have a median survival of 11 months ^33^. In solid organ transplants, a substantial proportion of patients experience mortality related to immune-mediated rejection and fibrosis ^34–36^. However, most immunosuppressive therapies used to prevent rejection broadly target the immune system, resulting in significantly increased infection risks ^37,38^. Therefore, therapies that specifically target subsets of immune cells such as T cells offer the potential for more precise control of the balance between infection risk and graft rejection.

In recent years, our understanding of T cell activation has expanded vastly with the advent of genome-wide screening^39^. Approaches using CD3 and CD28 antibodies to induce uniform T cell activation have significantly advanced our knowledge of TCR signaling ^40–42^. To study regulators of T cell function, direct engagement of T cells with their target cells are often required to uncover their functions. Thus, others have studied the direct effects of gene deletion or activation on T cell cytotoxicity *in vitro* and *in vivo* with cancer and viral models^43–48^.

Since large-scale analysis of T cell cytotoxicity can be challenging, an alternative strategy involves identifying proteins proximal to T cell signaling complexes during activation, with the assumption that spatial proximity correlates with functional significance ^49–51^. Next, more detailed cytotoxicity assays with T cells may be used.

To add to this body of work, we fused APEX2, a proximity labeling enzyme, to CD8ɑ, a coreceptor of the T cell receptor (TCR), to catalog the proteins that preferentially engages with it during T cell interaction with target cancer cells ^52^. This would in theory allow for labeling of possible regulators at the level of protein assembly, signaling, and degradation.

## Results

### CD8ɑ-APEX2 labels TMX1, an essential gene for T cell cytotoxicity

To catalog proteins around CD8ɑ during T cell activation, we used Jurkat T cells and melanoma line M257 and introduced the 1G4-TCR interaction with NY-ESO antigen. The target line M257 was edited to express HLA-A2 and pulsed with NY-ESO1 (SLLMWITQC) peptide and Jurkat T3.5 cells known to have no endogenous TCRβ and CD8 was edited to express 1G4 TCR and CD8ɑ-APEX2 (Fig 1A). Four hours after co-culture of those two cell lines, APEX2 was catalyzed with H_2_O_2_ and found to have higher biotin-labeling activity in co-cultures with NY-ESO-presenting cells compared to co-cultures with the B2m knockout negative control (Fig 1B). Intriguingly, CD8ɑ-APEX2 stabilization was found to be antigen dependent (*data not shown*). The labeled proteins were purified with streptavidin beads and characterized through mass spectrometry. We found that CD8ɑ-APEX2 labeled proteins were enriched for known T cell signaling proteins such as CD3δ, LCK, and ZAP70, supporting the validity to characterize the functional relevance of other hits (Fig 1C).

**Fig 1.**
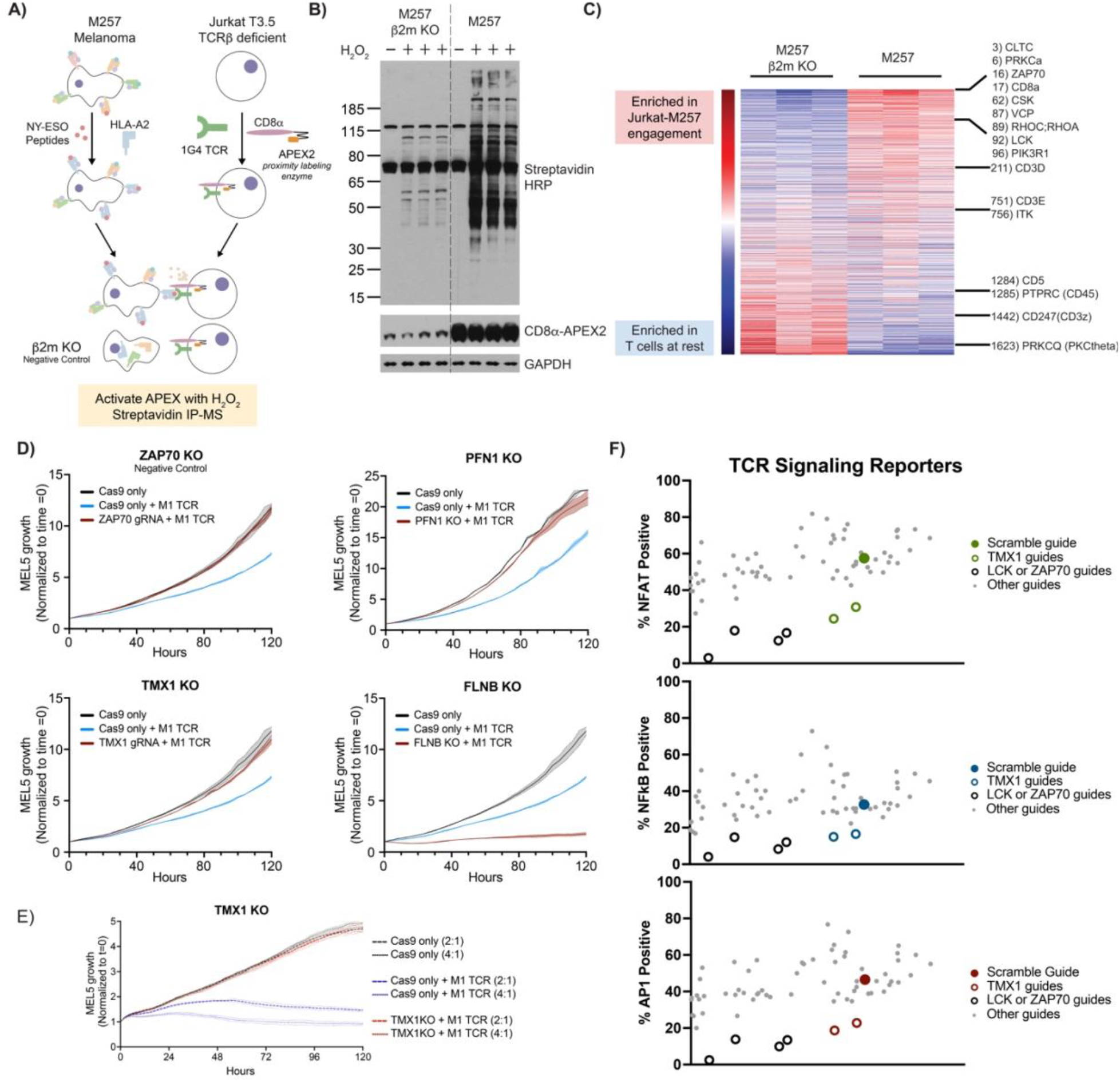
CD8ɑ-APEX2 labels TCR signaling proteins and identifies TMX1. **A)** Schematic of CD8ɑ-APEX2 labeling. **B)** Western blot of APEX labeled proteins 4 hours after co-culture between M257 and Jurkat with SA-HRP, CD8ɑ-APEX2, and GAPDH. **C)** Mass spectrometry result of CD8ɑ-APEX2 labeled proteins. **D)** MEL5 killing by M1 TCR expressing primary human CD8 T cell with individual genes deleted. **E)** MEL5 killing by different ratios of M1 TCR expressing primary human CD8 T cells with TMX1 deleted. **F)** NFAT, NFκB, and AP1 transcription activity in Jurkats with genes deleted activated by soluble aCD3aCD28.

We chose 32 hits and tested their impact on T cell cytotoxicity using primary human CD8 T cells isolated from PBMCs. For each hit, the T cells had that single gene deleted and transduced with the M1 TCR specific to the MART1 antigen. These modified CD8 T cells were then co-cultured with MEL5 melanoma cells expressing GFP and endogenous MART1. T cell cytotoxicity was measured using the Incucyte SX5 for decrease in GFP as a proxy for cancer cell death. This revealed that deletion of TMX1 or PFN1 significantly suppressed T cell cytotoxicity, whereas deletion of FLNB enhanced it (Fig 1D and 1E).

To investigate the pathways affected by these genes, we generated single cell clones of Jurkat E6.1 cells carrying NFAT, NFκB, and AP1 transcriptional reporters^53^, deleted each gene, and activated them with soluble αCD3αCD28 antibodies. TMX1 deletion was found to be deficient in all NFAT, NFκB, and AP1 (Fig 1F). Despite general consistency between the cytotoxicity and signaling reporter assays, the deletion of CSK resulted in increased NFAT, NFκB, and AP1 signaling although decrease in T cell cytotoxicity (*data not shown*).

### TMX1 is necessary for TCR signaling

TMX1 appeared to be a suitable target for immunosuppression since TMX1 deletion reduced T cell activity, TMX1 belongs to the oxidoreductase family which are druggable, and TMX1 deletion had been shown to be viable in mice ^54^. The known phenotypes of TMX1 deletion are increased platelet coagulation and increased susceptibility to liver damage^54,55^. Additionally, TMX1 is known as an ER resident transmembrane disulfide oxidoreductase that preferentially acts on transmembrane proteins ^56^. Single cell clones of TMX1-deleted Jurkat cells had significantly fewer cells with detectable NFAT, NFκB, or AP-1 activity when activated with soluble αCD3αCD28. Furthermore, prolonged incubation with the activation antibodies resulted in minimal increases in activity (Fig 2A). The deletion of TMX1 had little to no effect on the growth of the cells (Fig 2B), consistent with prior reports of TMX1 deletion mice were viable and without overt phenotypes. Lentiviral overexpression of TMX1 but not the catalytic inactive mutant of TMX1 (TMX1^*Inactive* (C56A|C59A)^) was able to rescue the signaling deficits (Fig 2C).

**Fig 2.**
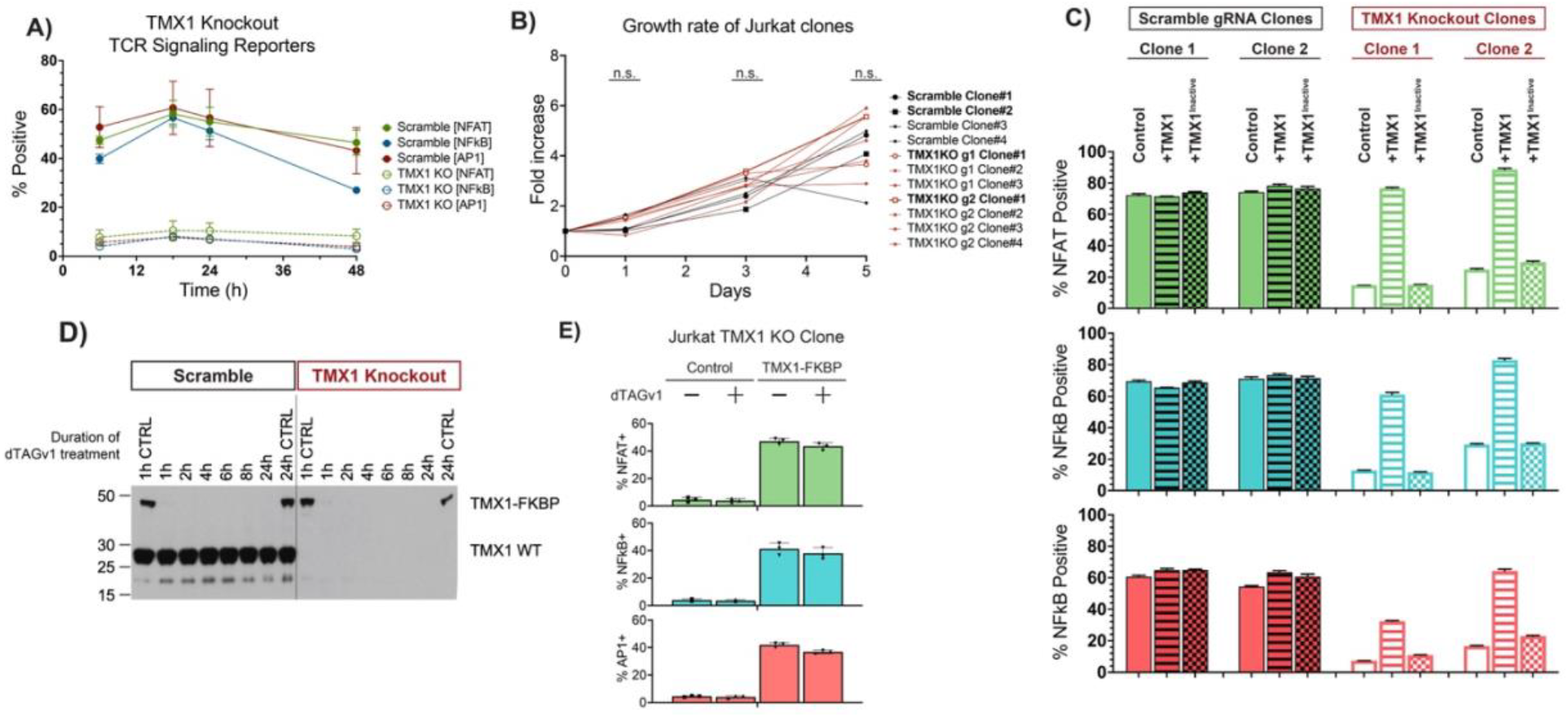
TMX1 catalytic activity is required for NFAT, NFκB, and AP1 signaling. **A**) NFAT, NFκB, and AP1 transcriptional activity in a single cell clone of TMX1 deleted Jurkat over time after stimulation with aCD3aCD28 antibodies. **B)** Growth rate of various clones of Jurkat with TMX1 deletion or with scramble gRNA. **C)** NFAT, NFκB, and AP1 transcriptional activity with overexpression of TMX1 or TMX1^*Inactive* (C56A|C59A)^. **D)** Western blot of TMX1 in Jurkat expressing TMX1-FKBP treated with dTAGv1. **E)** NFAT, NFκB, and AP1 transcriptional activity in Jurkat clone with TMX1 deletion expressing TMX1-FKBP, treated with dTAGv1, and activated with aCD3aCD28.

To test whether TMX1 was required during T cell activation, we generated TMX1 fused to FKBP12^F36V^ and used the dTAGv1 system to temporally control the degradation of TMX1^57^. Within 1h of dTAGv1, only trace levels of TMX1 were detectable by western blot (Fig 2D). When the cells were treated with dTAGv1 for 30 min followed by 6 hours of incubation with soluble αCD3αCD28, there appeared to be no changes in the NFAT, NFκB, and AP1 signaling (Fig 2E), suggesting TMX1 is not required during activation.

### TMX1 deletion causes decrease in CD3ζ and surface TCR

Since TMX1 deletion led to deficits in all three NFAT, NFκB, and AP1 signaling pathways, the mechanism is likely related to the early events of T cell signaling. To test this, we checked protein levels of common T cell signaling proteins in TMX1 deleted Jurkat clones. The western blot of TMX1 Jurkat clones revealed a strikingly specific deficit in CD3ζ (Fig 3A). There was a corresponding decrease in surface TCR and CD3 level as measured by flow cytometry (Fig 3B), consistent with prior studies of CD3ζ deletion leading to low surface TCR levels ^58,59^. However, CD3ζ mRNA level was not affected by TMX1 deletion (Fig 3C), suggesting the decrease in CD3ζ occurs at the post-transcriptional level. This deficiency in surface TCR could be rescued by lentiviral overexpression of TMX1 but not TMX1^*Inactive* (C56A|C59A)^ (Fig 3B). These results were also mirrored in primary human T cells, where TMX1 deletion led to a similar decrease in surface TCR and CD3 level (Fig 3D and 3E).

**Fig 3.**
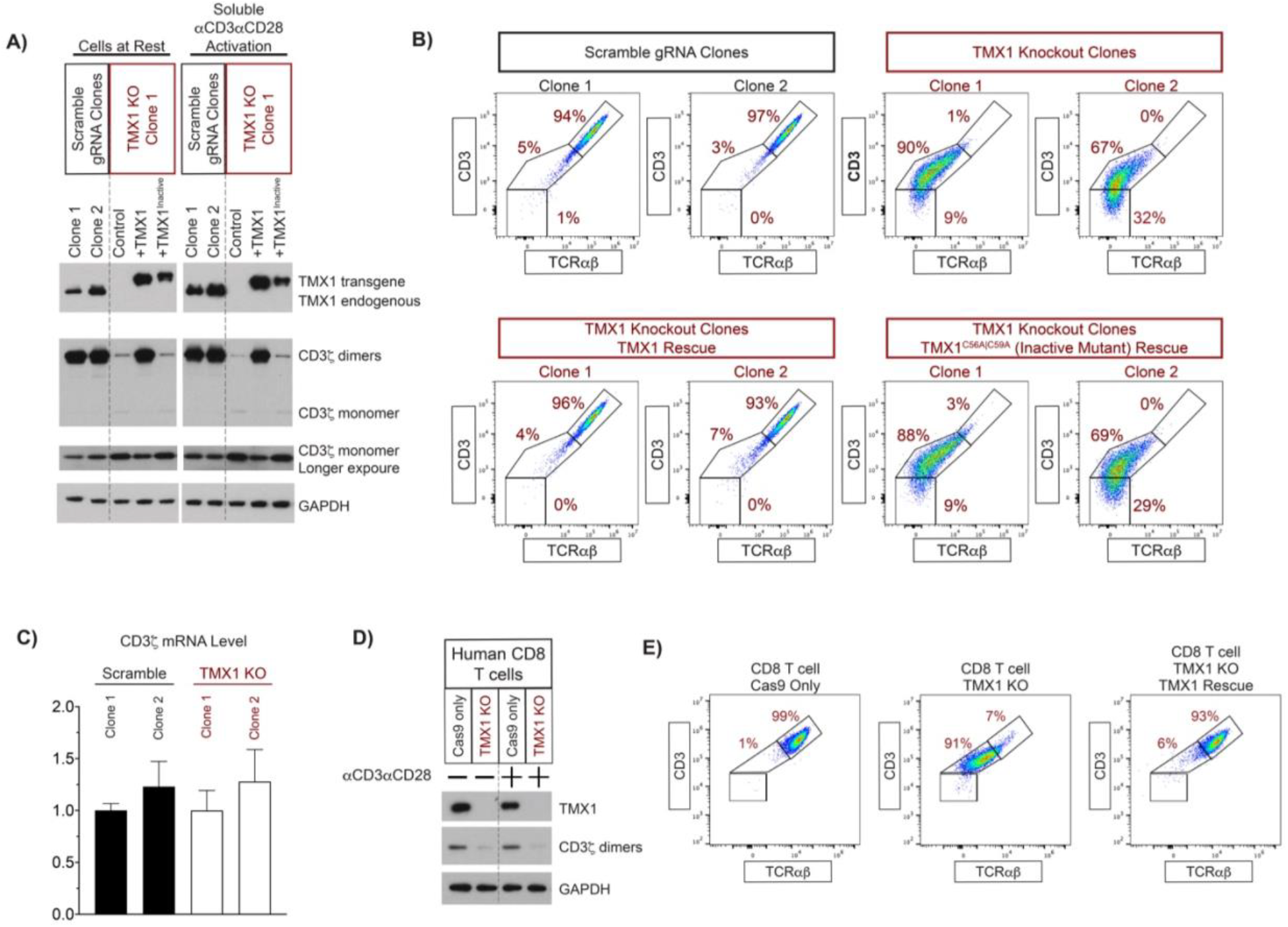
TMX1 deletion causes decrease in CD3ζ and surface TCR. **A)** Western blot of TMX1 and CD3ζ in Jurkat clones with TMX1 deletion overexpressing TMX1 or TMX1^*Inactive* (C56A|C59A)^. **B)** Flow cytometry of CD3 and TCRɑβ in Jurkat clones with TMX1 deletion overexpressing TMX1 or TMX1^*Inactive* (C56A|C59A)^. **C)** qPCR of CD3ζ mRNA in Jurkat clones with TMX1 deletion. **D)** Western blot of TMX1 and CD3ζ in primary human CD8 T cells with TMX1 deletion. **E)** Flow cytometry of CD3 and TCRɑβ in primary human CD8 T cells with TMX1 deletion with TMX1 overexpression.

### TMX1 deletion is mediated by downstream CD3ζ downregulation

To test whether TMX1 is necessary for CD3ζ function, CD3ζ was overexpressed in TMX1 knockout clones. We found that CD3ζ was sufficient to rescue the surface TCR levels in TMX1 deleted Jurkats (Fig 4A). Furthermore, CD3ζ overexpression was able to rescue the NFAT, NFκB, and AP1 signaling defects in these cells (Fig 4B).

**Fig 4.**
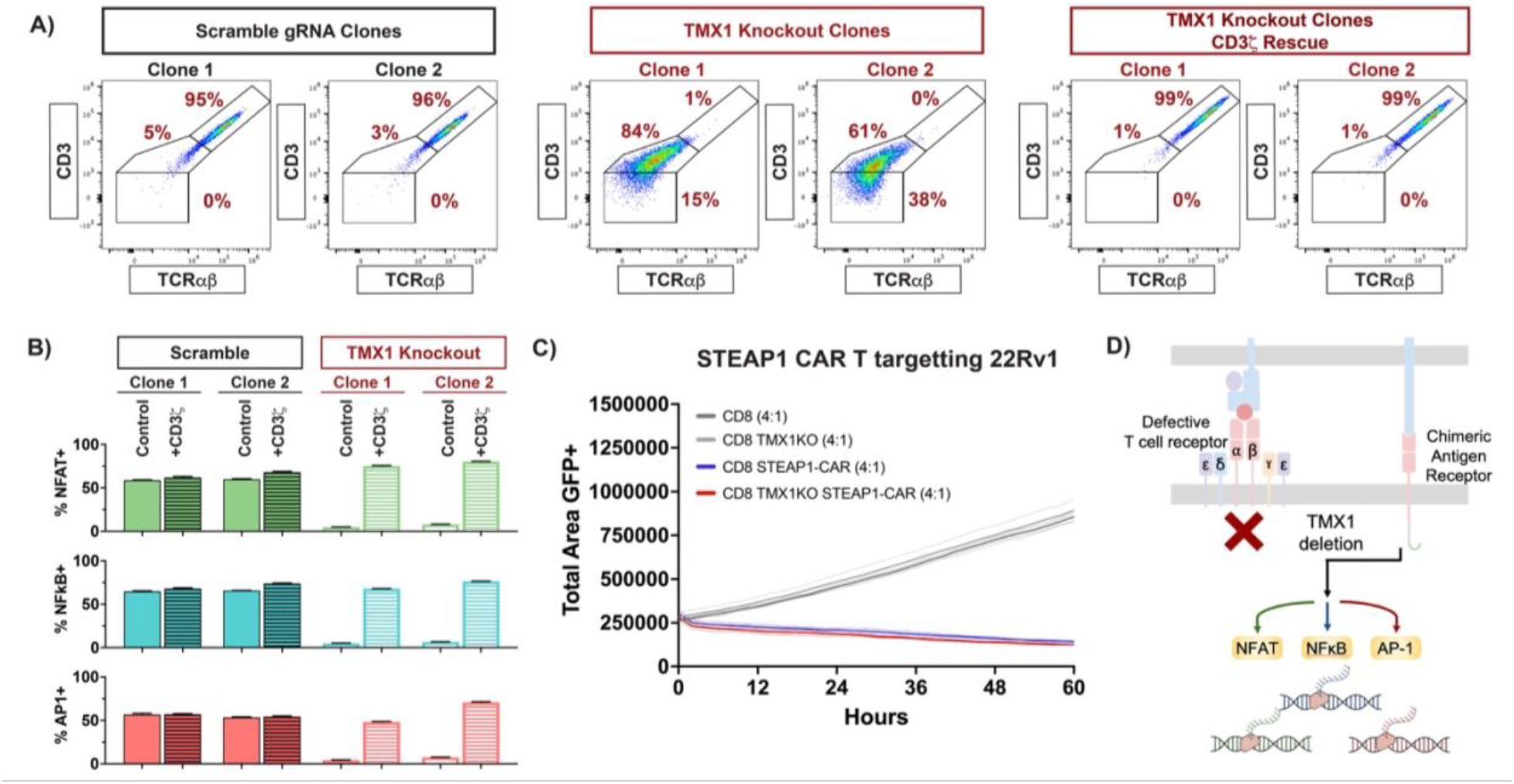
CD3ζ overexpression and CAR rescues T cell dysfunction in TMX1 deletion. **A)** Flow cytometry of CD3 and TCRɑβ in primary human CD8 T cells with TMX1 deletion overexpressing CD3ζ. **B)** NFAT, NFκB, and AP1 transcriptional activity in a single cell clone of TMX1 deleted Jurkat overexpressing CD3ζ after stimulation with aCD3aCD28 antibodies. **C)** 22Rv1 killing by STEAP1-CAR T with TMX1 deleted. **D)** Schematic of CAR T bypassing defective TCRs.

To show that CD3ζ downregulation was the only factor impacting T cell cytotoxicity, we transduced PBMC derived T cells with a STEAP1 targeting chimeric antigen receptor (CAR)^60^. Since the STEAP1 CAR carries its own CD3ζ, endogenous levels of CD3ζ or TCR assembly should not impact its function (Fig 4D). Using 22Rv1 (STEAP1+) cells with GFP, 22Rv1 with STEAP1 knockout, and 22Rv1-STEAP1 knockout with STEAP1 rescue, we found that TMX1 deletion has no impact on T cell cytotoxicity (Fig 4C). Together, these results suggest that the T cell dysfunction is solely caused by decrease in CD3ζ which is likely a downstream mechanism of TMX1 deletion.

### CD3δ is a substrate of TMX1

To identify the cause of CD3ζ destabilization due to TMX1 deletion, we searched for TMX1 substrates. Since the substrate interactions with disulfide oxidoreductases are often short lived, a substrate trap form of TMX1^*Trap* (C59A)^ is needed. As previously described^61^, C59 of TMX1 is required to break the disulfide intermediate between TMX1 and its substrate, mutation to this residue leads to a covalent trapping and stabilization of TMX1-substrate dimer (Fig 5A). We introduced Flag-tagged TMX1, TMX1^*Trap* (C59A)^, and TMX1^Inactive (C56A|C59A)^ separately into parental Jurkat E6.1 cells. The cells were either cultured for 24h with or without soluble aCD3aCD28 antibodies. Non-reducing western blot for TMX1 identifies many high molecular weight bands that can be eliminated with the addition of DTT (Fig 5B), indicating the bands represent TMX1 fused to its substrates. Flag tag magnetic bead IP was done to purify TMX1 and its substrates, followed by release of the substrates for mass spectrometry. CD3δ was found to be enriched in the samples with substrate trap TMX1^*Trap* (C59A)^ when compared to that of TMX1^Inactive (C56A|C59A)^ (Fig 5C).

**Fig 5.**
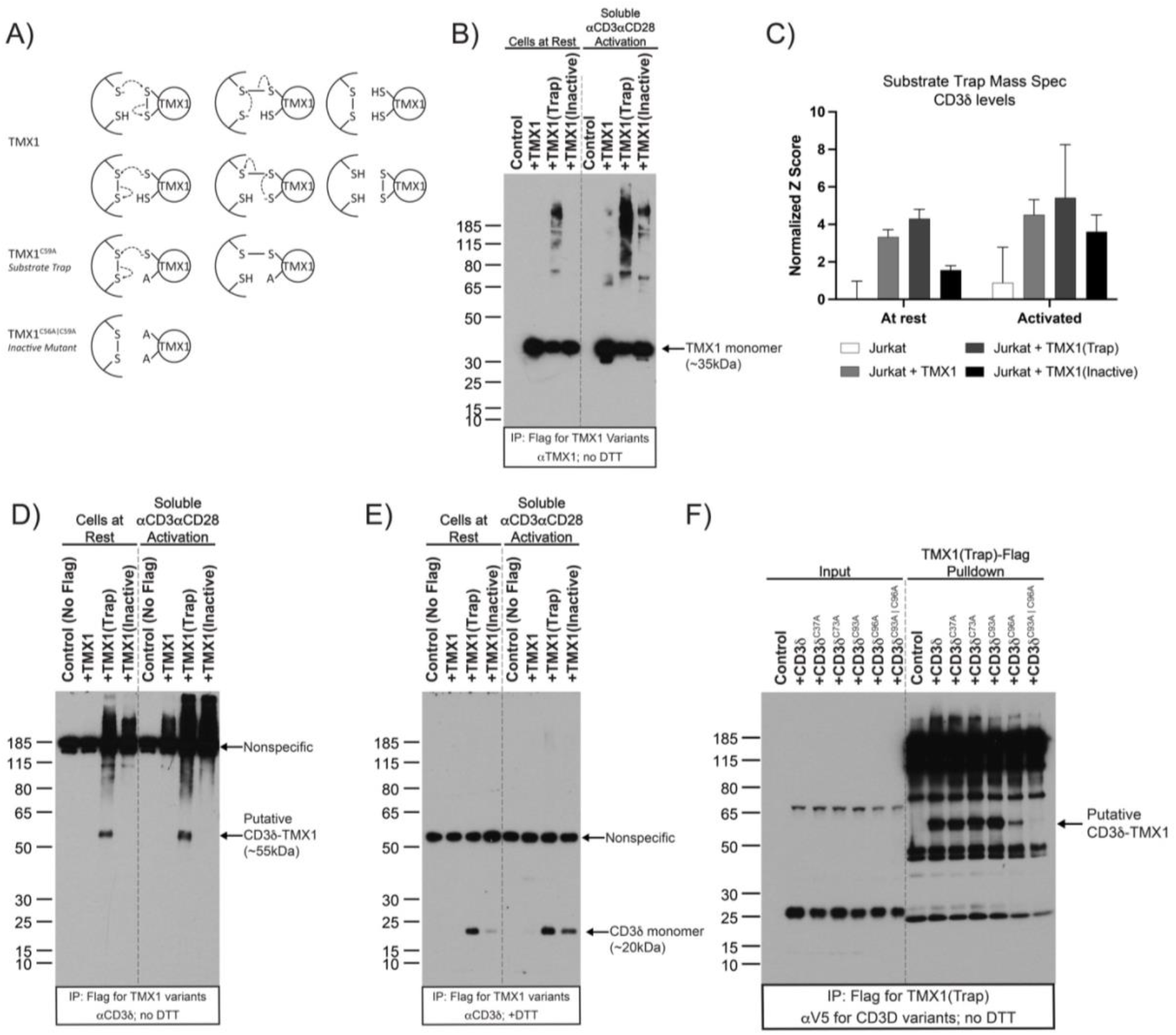
TMX1 forms disulfide bond with CxxC motif in CD3δ. **A)** Schematic of different forms of TMX1 mutants and its expected mechanisms. **B)** Western blot without DTT for TMX1 in flag tag IP of Jurkat expressing flag tagged TMX1, TMX1^*Trap* (C59A)^, or TMX1^Inactive (C56A|C59A)^. **C)** CD3δ level from mass spectrometry of TMX1 substrates. **D)** Western blot without DTT for CD3δ in flag tag IP of Jurkat expressing flag tagged TMX1, TMX1^*Trap* (C59A)^, or TMX1^Inactive (C56A|C59A)^. **E)** Same as D except with DTT added. **F)** Western blot without DTT for V5 tagged CD3δ in flag tag IP of Jurkat expressing flag tagged TMX1^*Trap* (C59A)^ and V5 tagged variant of CD3δ.

To validate this, TMX1 Flag-tag IP was done and blotted for CD3δ which showed a band at molecular weight of ∼55kDa and a smear from 80 to >180kDa (Fig 5D). The band seen at 185kDa was likely artifact from the IP process since it is present even in the control samples without any Flag-tagged proteins. The smear from 80 to >180kDa is likely to represent CD3δ coupled with various components of TCR subunits (Fig 5D). When DTT was added to remove all TMX1-substrate covalent bonds, CD3δ was found at ∼20kDa (Fig 5E) and a nonspecific band observed at ∼55kDa. Since TMX1 monomer was ∼35kDa in Fig 5B and CD3δ monomer was ∼20kDa in Fig 5E, this further strengthens the argument that the observed band at ∼55kDa in the non-reducing blot in Fig 5D likely represents the covalently bonded TMX1-CD3δ.

Notably, this IP enriched for all TMX1 associated proteins including substrates which differed from the substrate trap mass spectrometry experiment described in Fig 5C which enriched for just TMX1 substrates. With that in mind, in the samples with TMX1^*Inactive* (C56A|C59A)^, CD3δ was observed in the blot containing DTT as monomer at ∼20kDa (Fig 5E) though not as the putative ∼55kDa TMX1-CD3δ dimer (Fig 5D). Furthermore, T cell activation with aCD3aCD28 increased association of TMX1 with TCR complex as demonstrated by increased smear from 80kDa to 150kDa (Fig 5D). These results suggest an interaction between TMX1 and CD3δ which does not require the cysteine covalent bonds.

To identify the residues by which CD3δ interacts with TMX1, we generated point mutations to cysteine residues found in CD3δ localized within the ER lumen where TMX1 activity is found. Next, CD3δ cysteine mutants were introduced into Jurkat expressing TMX1^*Trap* (C59A)^. Through this, we found that C93 and C96 was required for the TMX1-CD3δ covalent interaction formation (Fig 5F). Notably, previously works have shown that this CxxC motif on CD3δ must be oxidized for proper T cell function^62,63^.

## Discussion

To identify new targets of T cell immunosuppression, we cataloged the interactome of CD8ɑ during T cell activation. Through this, we found that TMX1 is critical for TCR assembly and T cell function. Specifically, CD3ζ levels decreased significantly with TMX1 deletion. CD3ζ overexpression was sufficient to rescue the phenotype and CAR-T function was not impacted by TMX1 deletion. TMX1 was found to directly interact with the CxxC motif of CD3δ. These findings suggest that TMX1-mediated redox regulation of CD3δ may be essential for efficient TCR assembly (Fig 6). These results highlight the potential of targeting T cell receptor assembly for potent inhibition of T cell functions.

**Fig 6.**
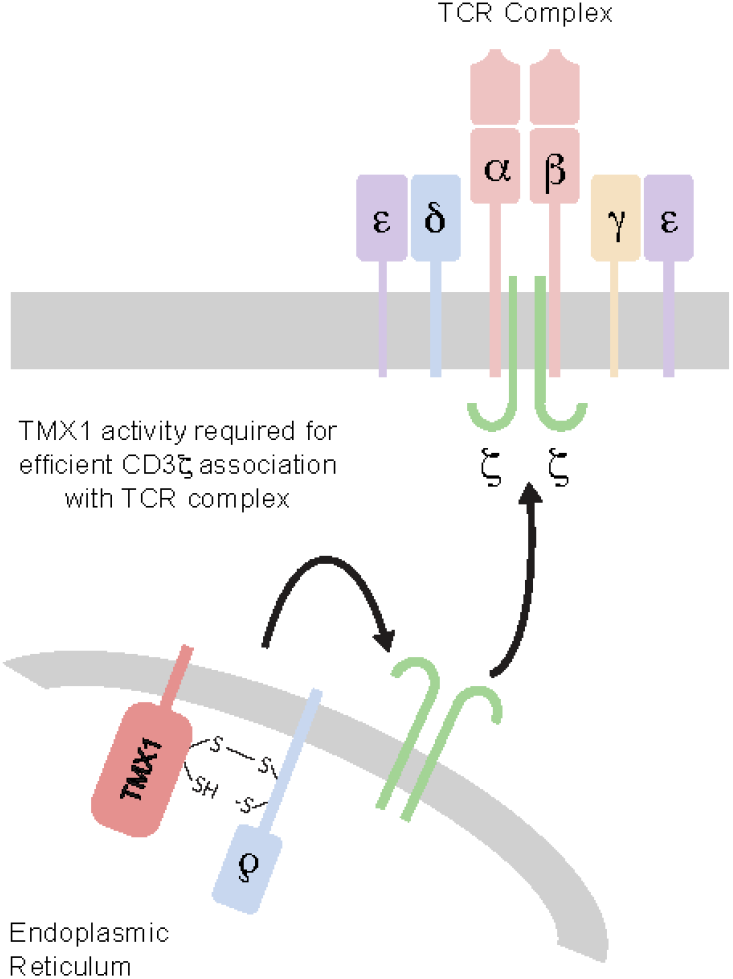
Graphical summary of TMX1’s impact on T cells.

Regulation of the TCR and particularly CD3ζ appears to be a critical node of T cell regulation involved in cancers and autoimmune disorders^64^. Previous work has shown that CD3ζ is rapidly degraded and is the rate limiting step to TCR assembly ^65,66^. CD3ζ has been described to be regulated through a variety of pathways including ROS^67^, T cell activation^68^, HSP10^69^, arginine ^70–72^, and GCN2^73^ ; however, we did not observe direct links between these pathways and TMX1 deletion (*data not shown*). Despite the prevalence of CD3ζ downregulation in pathologies, there are unsolved questions regarding the mechanisms.

In TMX1 deletion, we hypothesize that the decrease in CD3ζ stability is due to improper TCR assembly, CxxC motif of CD3δ is constitutively oxidized and required for interactions with TCRɑ ^62,63^. CD3δ deletion is also known to cause loss of surface TCR^74^, and cysteine to serine SxxS mutant of CD3δ led to T cell deficiency though no impact on surface TCR expression ^63^. This was consistent with our findings that CD3δ or CD3δ AxxA failed to rescue the phenotype observed in TMX1 deletion (*data not shown*). While we provide evidence that TMX1 engages the CxxC motif of CD3δ, the precise effect of TMX1 on its redox state remains unclear. The TMX1^*Trap* (C59A)^ we used is likely only able to identify substrates undergoing reduction but not oxidation. Thus, we propose that TMX1 could interact with CD3δ by: 1) reducing the oxidized CxxC bond on CD3δ; 2) oxidizing the reduced CxxC bond on CD3δ facilitating disulfide isomerization of another oxidized disulfide bond, potentially resulting in the formation of an oxidized CxxC on CD3δ; or 3) mediating a complex redox exchange involving multiple proteins ultimately leading to oxidation of CxxC on CD3δ. However, we are unaware of any method to express a constitutively oxidized form of CD3δ to prove test this hypothesis. Alternatively, TMX1 may regulate CD3ζ independent of CD3δ and act on a substrate which we could not detect.

Although further investigation is required to clarify how TMX1 deletion leads to CD3ζ downregulation, TMX1 deletion has a substantial impact on CD3ζ stability and T cell function. To our knowledge, no current therapies target the production of T cell receptors. Current treatments either prevent downstream TCR signaling or cause lymphopenia which is not acutely reversible in cases of infections. Thus, targeting TCR assembly and CD3ζ stability, which is not required for maintenance of existing T cells but inhibits their function may provide a new class of treatment option. In terms of feasibility, many medications are used to target oxidoreductases (e.g., allopurinol, methotrexate, atovaquone, monoamine oxidase inhibitors). Therefore, it may be possible to develop a TMX1 inhibitor. Proteins assembly chaperones are often common essential gene and expected to have significant side effects if inhibited. Surprisingly, TMX1-deficient mice did not display any overt abnormalities and T cells had no dysfunction in proliferation in our hands ^54^. However, since TMX1 is known to negatively regulate coagulation, therapies targeting it may increase the risk of hypercoagulability ^54^. Nonetheless, there may exist a therapeutic window for which TMX1 inhibition leads to T cell inhibition without causing significant side effects. Exploring TMX1 and other proteins that regulate the assembly T cell receptor complex may present a novel approach to treat conditions caused by T cell hyperactivity.

## Acknowledgements

T.C. is supported by the Stanford Medical Scientist Training Program (NIH T32GM007365). This work was supported by NIH/NCI Outstanding Investigator Award R35-CA220434 to I.L.W., the Virginia and D.K. Ludwig Fund for Cancer Research to I.L.W.

## Methods

### Cell line maintenance

M257(UCLA), Jurkat Clone T3.5(ATCC TIB-153), Jurkat Clone E6-1 (TIB-152), MEL-5 (ATCC HTB-70) were cultured in RPMI + Glutamax supplemented with 10% FBS and 1x Penicillin-Streptomycin. HEK293T/17 (ATCC ACS-4500) and A375 (ATCC CRL-1619) was cultured in DMEM + Glutamax supplemented with 10% FBS and 1x Penicillin-Streptomycin

### β2m Deletion in M257

pSpCas9(BB)-2A-GFP (PX458) was a gift from Feng Zhang (Addgene plasmid # 48138 ; http://n2t.net/addgene:48138 ; RRID:Addgene_48138). SgRNA targeting B2m (CGTGAGTAAACCTGAATCTT) was cloned into pX458. FugeneHD (Promega Cat#E2311) was used to transfect the pX458 with B2m sgRNA into M257 as described by the manufacturer. Knockout cells were bulk sorted twice for B2m (2M2) and HLA-ABC (W6/32) negative using BD FACSAria II.

### 1G4 and CD8ɑ-APEX2 expression in Jurkat T3.5

Jurkat Clone T3.5 was transduced with pCCL-c-MNDU3 driving the expression of the 1G4 TCR. Subsequently, they were transduced with lentivirus carrying pHAGE6-CMV-CD8ɑ-3xG4S Linker-APEX2-IRES-GFP. GFP positive cells were sorted on BD FACSAria II. pCCL-c-MNDU3-X was a gift from Donald Kohn (Addgene plasmid # 81071 ; http://n2t.net/addgene:81071 ; RRID:Addgene_81071). pHAGE6 was a gift from David Baltimore.

### Triple Reporter Jurkat Activity Assays

Jurkat Clone E6-1 was transduced with retroviruses carrying fluorescent reporters of NFAT, NFκB, and AP1 transcriptional activity developed by *Jutz et al* ^53^. Single cell clones were sorted and expanded. For activation, 2ug/mL of UltraLEAF anti-CD3 and 4ug/mL of UltraLEAF anti-CD28 were added. Unless otherwise indicated, 6 hours after activation, the cells were read on flow cytometry for NFAT, NFκB, and AP-1 activity.

### Lentivirus production protocol

293T/17 [ATCC Cat# CRL-11268] are maintained below 80% confluency in DMEM + Glutamax supplemented with 10% FBS and 1x Penicillin and Streptomycin. Plates used for virus production are rinsed briefly with poly-L-Lysine (R&D Systems Cat#3438-200-01). 293T/17 are seeded at 110k/cm^2^. Around 24 hours later, at 80-90% confluency, transfection of plasmids was done with FugeneHD. pMDL (146ng/cm^2^), pVSVG (79.7ng/cm^2^), pREV (56.1ng/cm^2^), and third generation lentivirus plasmid (225ng/cm^2^ for every 10kb) were mixed to a final volume of 13.27uL/cm^2^ with OptiMEM. 1.194uL/cm^2^ of FugeneHD was added to the DNA mix, mixed by pipetting, and incubated for around 15 minutes at room temperature. Transfection mix was added drop-wise to cells without a media change.

16 to 20 hours after transfection, media containing transfecting reagents was aspirated and pre-warmed collection media was added at half volume (e.g., 1mL for 6 well plates and 5mL for 10cm plates). When concentration was not needed, DMEM with 10% FBS and 1x Penicillin and Streptomycin was used. When concentration of virus was needed, UltraCULTURE without Glutamine (Lonza Cat#12-725F) or AIM-V (cat#) with 1x Glutamax and 20mM HEPES was used. Next, Amicon Ultracel-100 columns were used to concentrate the virus following manufacturer protocol.

Viruses were stored in -80^°^C indefinitely and used with freeze-thaw unless a specific MOI was needed for the experiment.

### APEX Labeling of Jurkat and M257 Coculture

M257 were seeded in 6-well plates at 2.1875 × 10^6^ cells / well (230k cells/cm^2^) and incubated for 2 hours 20uM of NYESO peptide SLLMWITQC (ElimBio). Cells were rinsed two times with PBS. Jurkat T3.5 with or without CD8ɑ-3xG4S Linker-APEX2 are added at 1.09375 × 10^6^ cells / well in RPMI + Glutamax supplemented with 10% FBS and 1x Penicillin-Streptomycin. Unless otherwise indicated, 3 hours after co-culture in incubator, 0.5mM of biotin tyramide in prewarmed (RPMI + Glutamax supplemented with 10% FBS and 1x Penicillin-Streptomycin) was supplemented and plates were returned to incubator for an additional hour.

*Labeling* The labeling protocol is described in *Lam et al 2014* ^52^. Briefly, 1mM of H_2_O_2_ in (RPMI + Glutamax supplemented with 10% FBS and 1x Penicillin-Streptomycin) was supplemented to the wells to pulse label for 2 minutes at room temperature.

Subsequently, 2x cold quenching solution (20mM sodium ascorbate, 20mM sodium azide, 10mM Trolox) was added to equal parts of existing volume. Cells were kept cold on ice from this point forward. The wells were triturated and transferred to a conical tube and spun at 400g at 4 degrees for 5 minutes. The samples were subsequently washed with 1x cold quenching solution (10mM sodium ascorbate, 10mM sodium azide, 5mM Trolox) followed by a cold PBS wash. In experiments with more cells (e.g., mass spectrometry experiments), 10mL of 1x cold quenching solution was used. In other experiments, 1mL of cold 1x quenching solution was typically used.

Cells were lysed in RIPA lysis buffer containing protease inhibitor cocktail (Sigma) and PMSF. The samples were sonicated. Debris was pelleted by spinning the sample at 21,000g for 15 minutes at 4 degrees. Supernatant was stored at -80.

### PBMC-derived T cell culturing

T cell media consisted of AIM-V (ThermoFisher Cat#12055091) with 5% heat-inactivated human AB serum (Sigma Aldrich Cat# H3667-100mL), 1x Glutamax (ThermoFisher Cat# 35050061), 1x Beta-mercaptoethanol (Gibco Cat#21985023), and additional cytokines depending on the T cell subsets being cultured.

Culturing CD8 T cell required 5ng/mL (10.38IU/mL) of recombinant IL-2 (Peprotech 200-02) and 0.5ng/mL (14.35IU/mL) of recombinant IL-15 (Peprotech 200-15).

Culturing CD4 T cell required 5ng/mL (585IU/mL) of recombinant IL-7 (Peprotech 200-07) and 0.5ng/mL (14.35IU/mL) of recombinant IL-15 (Peprotech 200-15).

IL2 was reconstituted in 0.1%BSA and 100mM Acetic Acid in UltraPure Water at 100ug/mL. IL-7 was reconstituted in 0.1%BSA in UltraPure Water at 10ug/mL.

IL-15 was reconstituted in 0.1%BSA in UltraPure Water at 10ug/mL.PBMC-derived T cells were thawed and cultured in 1 × 10^6^ cells /mL of media containing 1:1 ratio of Dynabeads™ Human T-Activator CD3/CD28 for T Cell Expansion and Activation (ThermoFisher Cat# 11131D) in respective media. Cells were expanded daily by doubling the culture volume with fresh media roughly maintaining the culture at 1-2 × 10^6^ cells / mL.

### Nucleofection of PBMC T cell

Nucleofection was done as previously described except for changes listed below^75^. Briefly, 3-6 days after thaw and activation with Dynabeads™ Human T-Activator CD3/CD28, PBMC T cells were collected and Dynabeads are removed with DynaMag-2. RNP was formed at 1:2.5 molar ratio of Cas9 (IDT Cat#1081059) to total sgRNA (Synthego CRISPRevolution sgRNA EZ Kit). For a 20uL nucleofection reaction, 96pmol of Cas9 was added to 240pmol of sgRNA in 4uL and incubated for 10min. Next, cells resuspended in 15uL P3+supplement (Lonza Cat #V4XP-3032) and 1uL of TE buffer was added to the 4uL of RNP and nucleofected with EH-100 on 4D-Nucleofector System (Lonza).

The guides used for TMX1 knockout in primary human T cells were UAUUUCUUAGUUAUGCCCCG, GGUUGAAGAUUUUGACAAGC, and GUCAAAAUCUUCAACCGGAA.

### In vitro cytotoxicity assays

GFP+ cancer cells transduced with FUCGW are seeded at 2,500 cells per well of 96 well flat-bottom at the same time or one day before the addition of T cells. Cancer cells are counted the day after to determine the number of T cells needed to achieve 1:1 – 1:8 cancer to T cell ratio. T cells are resuspended in the cancer culture media (RPMI + GlutaMax + 10% FBS + 1x Penicillin-Streptomycin) before addition.

Incucyte SX5 is used to image once every hour to four hours with the 10x objective for 5 images per well. The Sartorius software was used to calculate surface area with GFP+.

### Substrate Trap Mass Spectrometry

TMX1, TMX1^*Trap* (C59A)^, and TMX1^Inactive (C56A|C59A)^ fused to flag tag followed by T2A EGFR^truncated^ were cloned into 3rd generation lentivirus plasmid pCCL-c-MNDU3. Jurkat E6.1 cell line (ATCC TIB-152) was transduced with lentiviruses carrying the plasmids above. The cells were cultured in RPMI + Glutamax supplemented with 10% FBS and 1x Penicillin-Streptomycin with or without anti-CD3 or anti-CD28 antibodies for 24 hours. Cells were collected and washed twice with cold 1x PBS containing 20mM NEM. Next, they were flash frozen with liquid nitrogen.

The cells were lysed in RIPA lysis buffer with 20mM NEM and protease inhibitor cocktail (Sigma). 80uL of anti-Flag M2 magnetic bead slurry (M8823-5ML) were used to pull down 2.3mg of lysate. The beads were rotating for 3 hours at 4 degrees followed by 5 washes with 1mL of RIPA lysis buffer without NEM or protease inhibitor.

For western blot analysis, 1/6th of the beads were resuspended in 60uL of 2x protein loading buffer without reducing agent. The samples were incubated for 10 minutes at 99 degrees and the supernatant was stored at -80.

For mass spectrometry, the remainder of the beads were resuspended in 80uL of 100mM TEAB. 3uL of 200mM TCEP were added and incubated at 65 degrees for 1 hour. After allowing samples to cool to room temperature for 5 minutes, 4.5uL of 200mM iodoacetamide resuspended in 100mM TEAB was added. The samples were incubated in the dark for 1.5 hours. 142.5uL of 100mM TEAB was added, and the supernatant was transferred to a new tube. 1mL of ice cold acetone was added and samples were allowed to precipitate overnight at -20.

Acetone precipitated samples were spun at 18,000g for 15 minutes at 4 degrees. Supernatant was removed carefully ensuring the protein pellet was not disturbed. The tubes were left open to let air dry. Protein pellets were resuspended in 100uL of 100mM TEAB and 5uL of Promega Sequencing Grade Trypsin (V5113).

Samples were incubated at 37 degrees for 18 hours. 41uL of anhydrous acetonitrile was used to dissolve each vial of the TMT10plex labeling reagent (Thermo Fisher Cat# 90309) and added to the digested samples. Labeling reaction occurred for 1 hour at room temperature. 8uL of 5% hydroxylamine was added to quench the reaction at room temperature for 15 minutes.

Equal amounts of the samples were mixed and stored at -20 for mass spectrometry.

## Notes

### Competing Interest Statement

The authors have declared no competing interest.

